# Validation of a small molecule inhibitor of PDE6D-RAS interaction with potent anti-leukemic effects

**DOI:** 10.1101/2022.03.14.484294

**Authors:** Sara Canovas Nunes, Serena De Vita, Andrew Anighoro, François Autelitano, Edward Beaumont, Pamela Klingbeil, Meaghan McGuinness, Beatrice Duvert, Chad Harris, Lu Yang, Sheela Pangeni Pokharel, Chun-Wei Chen, Monika Ermann, David A. Williams, Haiming Xu

## Abstract

RAS mutations prevalent in high-risk leukemia have been linked to relapse and chemotherapy resistance. Efforts to directly target RAS proteins have been largely unsuccessful. However, since RAS-mediated transformation is dependent on signaling through the RAS-related C3 botulinum toxin substrate (RAC) small GTPase, we hypothesized that targeting RAC may be an effective therapeutic approach in RAS mutated tumors. Here we describe multiple small molecules capable of inhibiting RAC activation in acute lymphoblastic leukemia cell lines. One of these, DW0254, also demonstrates promising anti-leukemic activity in RAS-mutated cells. Using chemical proteomics and biophysical methods, we identified the hydrophobic pocket of phosphodiester 6 subunit delta (PDE6D), a known RAS chaperone, as a target for this compound. Inhibition of RAS localization to the plasma membrane upon DW0254 treatment is associated with RAC inhibition through a phosphatidylinositol-3-kinase/AKT-dependent mechanism. Our findings provide new insights on the importance of PDE6D-mediated transport for RAS-dependent RAC activation and leukemic cell survival.

## Introduction

Guanosine triphosphatases (GTPases) are small G proteins that play key roles in hematopoietic cells in a variety of cellular processes including proliferation, apoptosis, cell migration, and cytoskeleton rearrangements [1] [2]. Activating mutations in RAS GTPase isoforms have been linked to numerous types of human cancers, including myeloid and lymphoid malignancies [3] [4] [5] [6]. NRAS/KRAS mutations have been found in 20%-25% of patients with acute myeloid leukemia (AML) [5], 25%-30% of patients with juvenile myelomonocytic leukemia (JMML) [7], and 15% of pediatric patients with B- or T-lineage acute lymphoblastic leukemia (ALL) [8] [9]. Specifically, RAS mutations are highly prevalent in relapsed high-risk ALL after combination chemotherapy, and the activation of RAS signaling has been shown to act as the driver of both *de novo* and relapsed, chemotherapy resistant disease [10] [11]. The various attempts to develop drugs that directly target mutant RAS proteins have been largely unsuccessful and to this day, only specific KRAS G12C inhibitors have been developed with evidence of clinical activity in solid tumors [12] [13]. However, this specific mutation is usually not found in relapsed acute leukemia patients [11].

Since, in some model systems, RAS-related C3 botulinum toxin substrate (RAC) GTPase is required for full RAS transformation [14] and leukemia cell survival [15] [16], we and others have focused on inhibiting its activity to indirectly target RAS signaling [17]. Here we report the identification of a compound DW0069 and development of two derivatives, DW0254 and DW0441, which demonstrated dose-dependent RAC inhibition, arrest of proliferation and induced apoptosis in human leukemic cell lines. We found that these compounds bind the hydrophobic pocket of phosphodiester 6 subunit delta (PDE6D), a RAS chaperone protein. Directed mutation of this pocket led to compound resistance, directly implicating molecule binding to PDE6D to cell growth inhibition. We further showed that treatment with DW0254 disrupts the interaction between PDE6D and RAS, disturbing RAS subcellular localization. Moreover, the dose-dependent decrease in RAC activation downstream of phosphatidylinositol 3-kinase/protein kinase B (PI3K/AKT) provides a biochemical link between RAS and RAC in leukemia cells. In summary, our study provides evidence that PDE6D-dependent RAS trafficking with downstream activation of PI3K/AKT and RAC constitutes a novel potential therapeutic target in high risk leukemias.

## Results

### Identification of small molecules demonstrating RAC inhibition

We hypothesized that targeting RAC could be an effective therapeutic approach for RAS-mutated leukemias.

Our initial screen for a RAC inhibitor depicted in Figure 1 lead to the identification of compound DW0069, and further medicinal chemistry efforts yielded the closely related compounds DW0254 and DW0441 (1, 2 and 3 respectively in Figure 2A) [18]. These early leads had suboptimal to satisfactory physiochemical properties although all showed improved biological activity on leukemia cells when compared to the tool RAC inhibitor NSC23766 [19] which showed cellular activities in the ∼40-80µM range ([20] and Figure 2A). In contrast, DW0346 analogue with an aliphatic amide substitution (4 in Figure 2A) showed a significant reduction in inhibitory activity on leukemic cells and was used as a negative control in subsequent target validation experiments. DW0254 was further profiled as it offered the best compromise between lipophilicity, solubility, and potent biological activity. DW0254 antileukemic activity was tested on a large panel of ALL and AML cell lines that exhibited varying levels of sensitivity to DW0254 (Figure 2B). 75% were considered responsive with a mean IC_50_ between 1 and 10µM. Treatment of P12-ICHIKAWA cells, with the lowest IC_50,_ caused a dose-dependent inhibition of RAC activation (Figure 2C), decrease in cell proliferation (Figure 2D) and increase in apoptosis (Figure 2E). DW0069 and DW0441 also affected cell growth, apoptosis and RAC activation (Supplemental Figure 1A-E). Unexpectedly, and in contrast with NSC23766, neither DW0069 or optimized DW0254 showed inhibition of the RAC1-TIAM1 protein-protein interaction as measured by homogeneous time-resolved fluorescence (HTRF) (Figure 2F and Supplemental Figure 1F).

**Figure 1.**
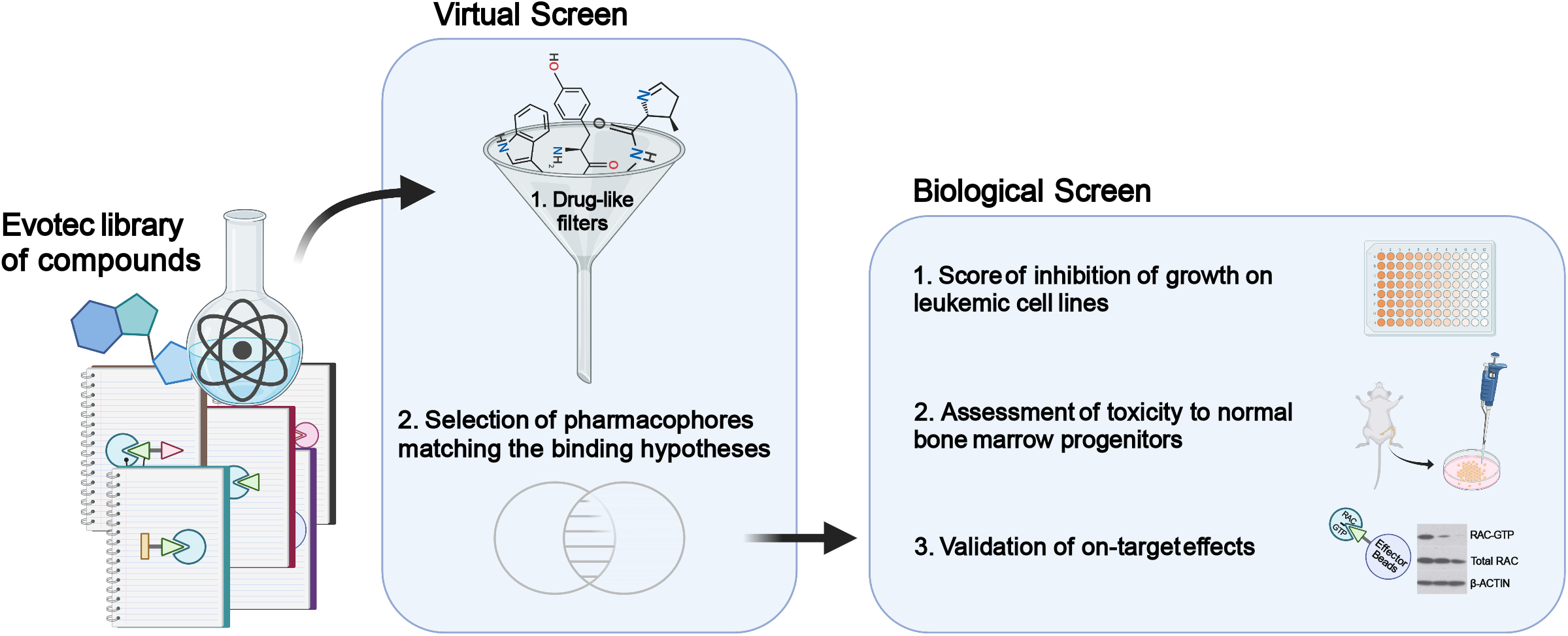
Compound screen for Rac inhibitors. A virtual screen of Evotec’s library of commercially available compounds was performed that included an initial filtration for drug-like properties, followed by a preselection against shape-based pharmacophore of published RAC inhibitors. Next, a biological screen was carried out consisting of: i) scoring for inhibition of growth on leukemic cell lines of 107 selected compounds, ii) assessment of toxicity to normal bone marrow progenitors of compounds that showed good anti-leukemic activity, and iii) validating on-target effects by RAC PBD pull down of non-toxic compounds. Created with BioRender.com.

**Figure 2.**
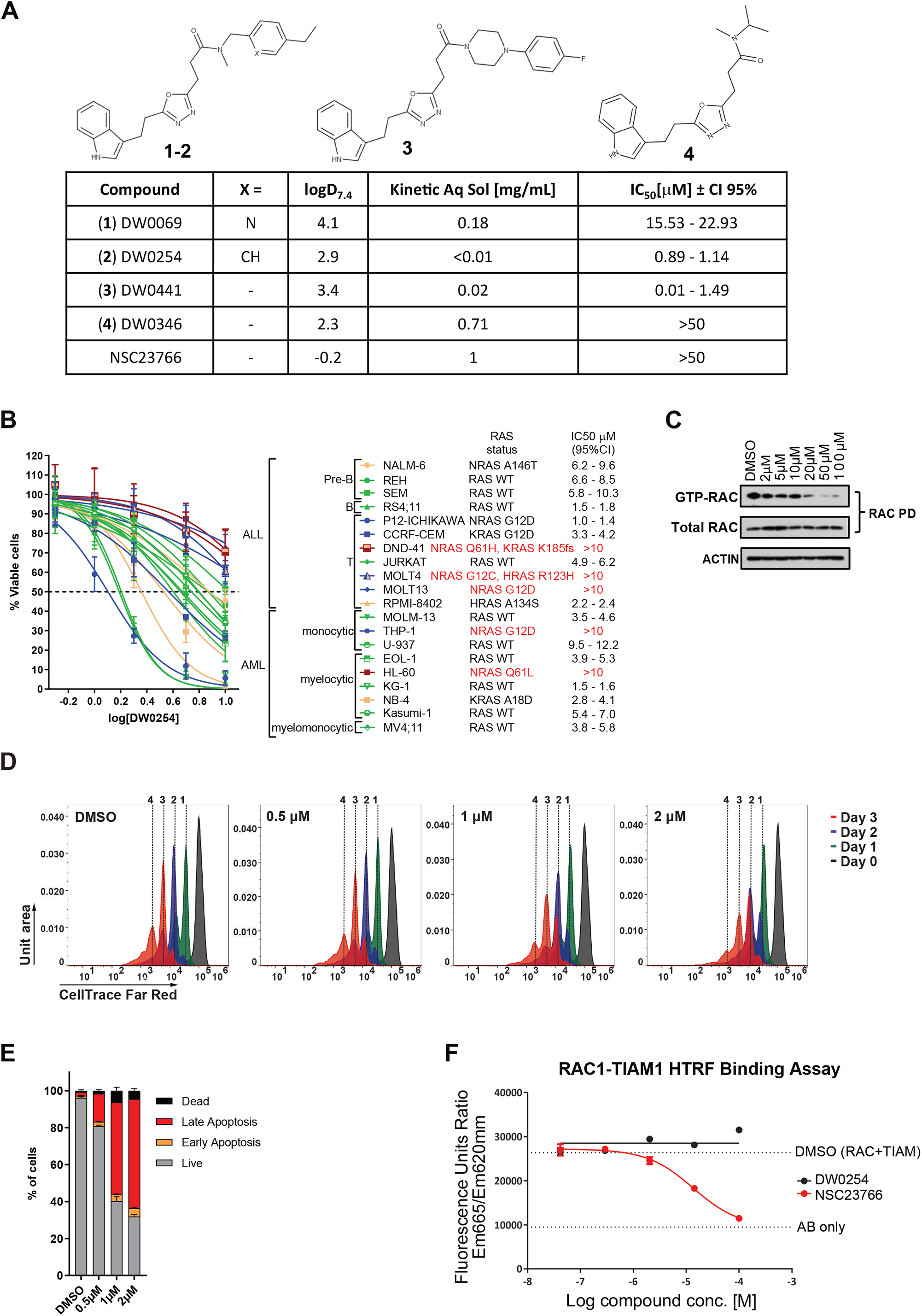
Compound DW0254 inhibits RAC activation and shows anti-leukemic activity in vitro in leukemia cell lines. **A)** Chemical structure, physicochemical properties, and biological activities cell line for compounds 1-4 and NSC23766 (structure not shown), a known inhibitor of RAC. IC_50_ values represent the dose at which 50% cell viability was achieved on P12-ICHIKAWA cells. **B)** Drug dosage curve showing live cell viability assay after 3 days of DW0254 treatment of human ALL and AML cell lines with diverse backgrounds and RAS status as described, n=4 at each dosage, data show mean ± SD, one of three individual experiments showing the consistent results. Color code: WT RAS green, G12 mutant RAS blue, Q61 mutant RAS burgundy, other RAS mutations yellow. **C)** GTP-RAC activity inhibition in P12-ICHIKAWA cells treated with different doses of DW0254 for 3 hours. GST pulldown assays were conducted by incubating lysates with PAK1-PBD beads. Cell lysates to detect total RAC and proteins eluted from the PAK1-PBD beads to detect GTP-RAC were subjected to Western blotting using anti-RAC (610651, BD Transduction laboratories, San Jose, CA) and anti-beta ACTIN (A5441, Sigma-Aldrich) antibodies. Data are representative of three individual experiments. **D)** Representative peaks of Far Red CellTrace staining of P12-ICHIKAWA cells treated with different doses of DW0254 and examined by FACS on three consecutive days. Peaks 1-4 represent the number of times the cells in each peak have divided; data shown from one of the three independent experiments. **E)** Bar graph showing percentage of apoptosis by AnnV/PI staining of P12-ICHIKAWA cells treated for 3 days with different doses of DW0254, data represent mean ± SD of 2 independent experiments with n=3 samples for each condition. Live: AnnV^-^/PI^-^; Early apoptosis: AnnV^+^/PI^−^; Late apoptosis: AnnV^+^/PI^+^; and Dead: AnnV^−^/PI^+^. **F)** Compound titration in the RAC1-TIAM1 homogeneous time-resolved fluorescence assay (HTRF) assay showing competition with increasing concentrations of test compounds (either NSC23766 or DW0254) on X-axis and fluorescence emission (Y-axis).

### Identification of PDE6D as target of DW0254

In contrast with the off-target effects exhibited by NSC23766, DW0069 chemical series showed no significant inhibition against a focused panel of kinases and G protein-coupled receptors (GPCRs) (Supplemental Figure 1G) [21]. With certain key pathway targets ruled out, we embarked upon on the deconvolution of the putative molecular targets of DW0254 using cellular photoaffinity labeling methods combined with label-free quantitative mass spectrometry analysis (PAL-MS). A PAL photoprobe consisting of the DW0254 warhead covalently linked to a minimalist terminal propargyl-diazirine photocrosslinker [22] was synthesized (Figure 3A). Like its parent compound, the PAL probe possessed antiproliferation properties (data not shown), demonstrating that the photoprobe was cell permeable and retained its activity.

**Figure 3.**
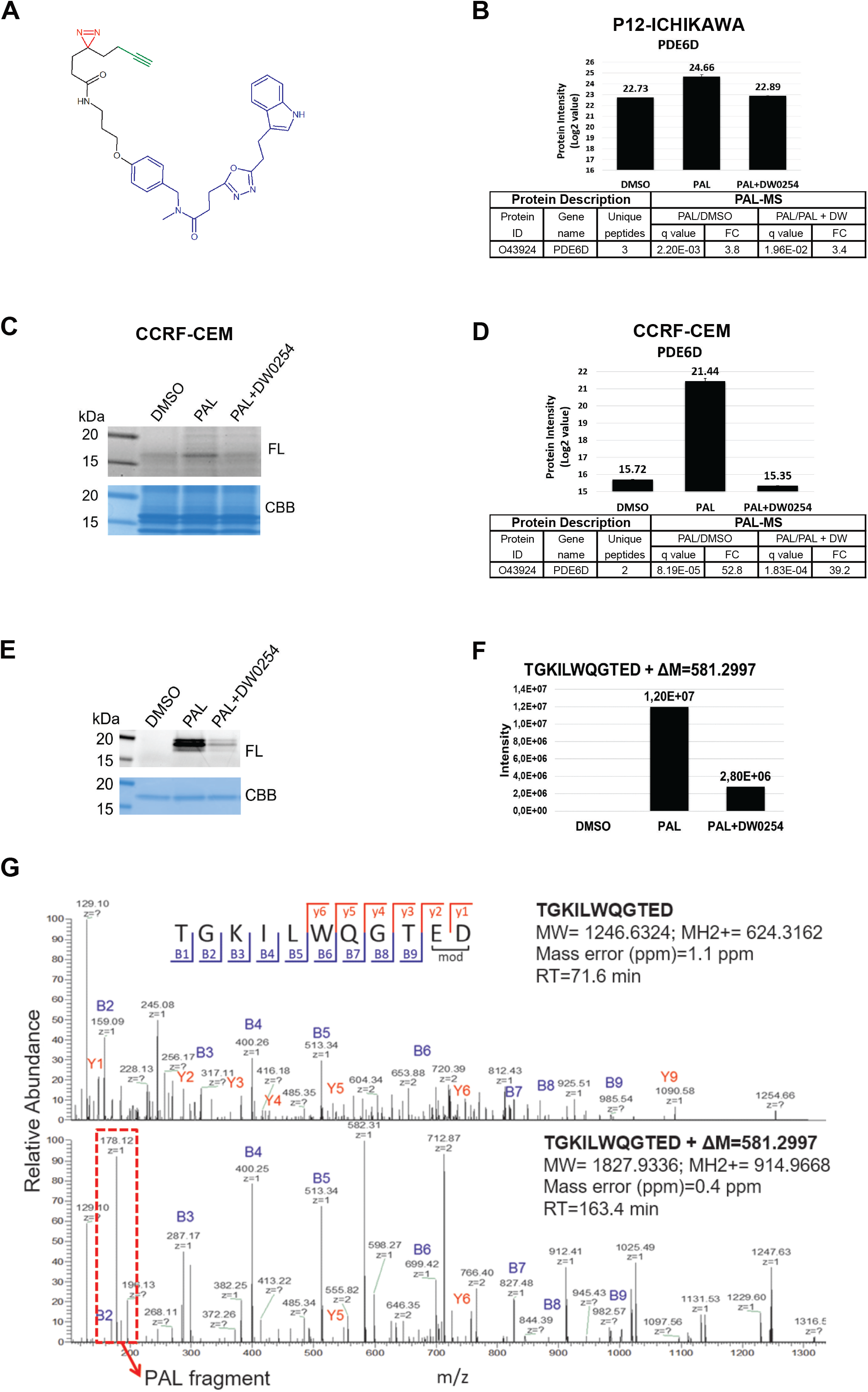
Identification of the DW0254 molecular target PDE6D by Photoaffinity Labeling Mass Spectrometry (PALMS). **A)** Chemical structure of DW0254-photoprobe PAL. The DW0254 warhead is colored in blue, the photoreactive diazirine group in red, and the alkyne clickable group in green. **B)** <u>top</u>: MS signal intensity of protein target hit of DW0254 in the pulldown samples of P12-ICHIKAWA cells. Histogram plots represent quantitative determination of PDE6D MS signal intensity in the different pulldown samples (DMSO, PAL alone, and PAL in combination with a 20-fold molar excess of DW0254). Conditions analyzed included P12-ICHIKAWA cells that were treated with PAL (1µM) ± DW0254 (20µM) prior UV irradiation, streptavidin pulldown and label-free differential quantitative mass spectrometry analysis. Three biological replicates for each sample were performed. <u>bottom</u>: Summary of the significant protein target hits of PAL identified in P12-ICHIKAWA; Proteins with an adjusted p-value (or q value) <5% and a FC of >2 were selected to be differentially modulated. A protein was considered as a hit of DW0254 when identified with at least two peptides in minimum 2 out of 3 replicates, FC>2 and adjusted p-values <0.05 in the two comparisons, PAL/DMSO and PAL/PAL+DW0254. **C)** In-gel fluorescence scanning showing the proteome reactivity profiles of live CCRF-CEM cells photolabeled by PAL (1µM) with or without DW0254 (20µM). FL = in-gel fluorescence scanning. CBB = Coomassie gel. **D)** <u>top</u>: MS signal intensity of PDE6D in CCRF-CEM pulldown samples. Histogram plots represent quantitative determination of PDE6D MS signal intensity in the different pulldown samples (DMSO, PAL alone, and PAL in combination with a 20-fold molar excess of DW0254). Y-axis shows log2 value of protein identified. Proteins eluted from the beads were separated by SDS-PAGE and protein bands within the molecular weight range 15-18kDa were excised. Proteins were prepared for downstream label-free differential quantitative mass spectrometry analysis. Three biological replicates for each sample were performed. <u>bottom</u>: Summary of the significant protein target hits of PAL identified in CCRF-CEM cells; Proteins with an adjusted p-value (or q value) <5% and a FC of >2 were selected to be differentially modulated. A protein was considered as a hit of DW0254 when identified with at least two peptides in minimum 2 out of 3 replicates, FC>2 and adjusted p-values <0.05 in the two comparisons, PAL/DMSO and PAL/PAL+DW0254. **E)** In-gel fluorescence scanning showing the recombinant human His-TEV-PDE6D-Avitag protein photolabeled by PAL (1µM) with or without DW0254 (20µM). **F)** Single-stage LC-MS (MS1) intensity values of PAL-modified peptide TGKILWQGTED following His-TEV-PDE6D-Avitag protein labeling with PAL in competition with DW0254. The peptide adduct was identified in the sample irradiated with PAL in the presence of DW0254 but with a peak intensity >4-fold lower compared to the sample irradiated with PAL alone. **G)** The second stage of mass spectrometry (MS2) for the PAL-modified peptide TGKILWQGTED of His-TEV-PDE6D-Avitag protein. MS2 spectra of the probe-modified peptide 1827.9216 m/z and its intact counterpart 1246.6193 m/z. Unlabeled fragment ions y1-y6 and b1-b9 were detected in both the PAL-modified peptide TGKILWQGTED and its intact counterpart. The fragment ion +178.12m/z cleaved from PAL1 upon CID fragmentation was detected only in the MS2 spectrum of the PAL1-modified peptide. FL: in-gel fluorescence scanning. CBB: Coomassie gel.

Retinal rhodopsin-sensitive cGMP 3’,5’-cyclic phosphodiester 6 subunit delta (PDE6D) was identified as a target hit in P12-ICHIKAWA cells with the highest signal intensity (Log2 Intensity) of 24.66 and with the highest sequence coverage of 28.6% (Figure 3B). We further confirmed PDE6D-PAL specific binding in an additional cell line, CCRF-CEM. Labeled protein patterns showed a protein band of ∼17kDa photolabeled with PAL probe that was protected by excess DW0254 (Figure 3C), further identified as PDE6D with a high signal intensity of 21.44 (Figure 3D). Photolabeling of recombinant human PDE6D expressed in *E*.*coli* also confirmed photoincorporation of the PAL probe into PDE6D that was fully protected by an excess of DW0254 (Figure 3E).

To gain insights into the binding site through identification of the specific photolabeled residues, recombinant PDE6D was UV-irradiated alone or with PAL probe in the presence or absence of DW0254 and analyzed by liquid chromatography–mass spectrometry (LC-MS/MS). A unique tryptic peptide of human PDE6D, TGKILWQGTED, was detected with an increase in peptide mass of +581.2997m/z corresponding to the incorporation of the PAL probe, with a >4-fold lower peak intensity in the presence of the competitor DW0254 (Figure 3F). PAL probe-modified peptide and its unlabeled control MS data were manually evaluated for the presence of specific probe-labeled b- or y-type fragment ions to further refine the localization of the photoadduct to a specific amino acid. Fragment ions y1-y6 and b1-b9 were detected in the unlabeled control TGKILWQGTED peptide (Figure 3G, top). The PAL-modified peptide (Figure 3G, bottom) shared the same fragment ions except for y1 and y2 suggesting that the PAL probe photolabeled, in a DW0254-inhibitable manner, residues E36 or D37 within the hydrophobic pocket of the molecule.

### Saturating mutagenesis screen hints at DW0254 binding mode

To further validate PDE6D as the biological target, to identify additional key residues for binding, and to link target engagement to the observed phenotype, we designed a sgRNA library (Supplemental Table 1) and performed a saturation mutagenesis screen of PDE6D. spCas9-expressing P12-ICHIKAWA cells were transduced with the PDE6D sgRNA library and treated with DW0254 with the goal of selecting resistant cells. After 2 weeks of treatment, 35% of library-transduced cells were alive, compared to 3% of the empty vector control cells (Figure 4A). A robust editing efficiency was confirmed by the decrease in positive control sgRNAs that targeted essential genes (Figure 4B). Specific sgRNAs were significantly enriched after DW0254 treatment, including sgRNA#144, which was identified at 20-fold increased frequency relative to DMSO-treated cells (Figure 4C, blue dot) and cells exhibited a ∼3-fold higher IC_50_ when compared to untreated library transduced cells (Figure 4D), confirming decreased compound sensitivity. Deltrasin, a commercially available PDE6D inhibitor, and additional derivative compounds have previously been reported to bind in PDE6D’s hydrophobic pocket and inhibit the growth of pancreatic cancer cell lines [23]. Since our data suggests the same binding site for DW0254, we tested DW0254-treated PDE6D-edited cells for sensitivity to Deltarasin. While unedited cells displayed a higher IC_50_ to Deltarasin (grey in Figure 4E) when compared to DW0254 (grey in Figure 4D), DW0254-treated PDE6D-edited cells demonstrated no increased resistance to Deltarasin (Figure 4E).

**Figure 4.**
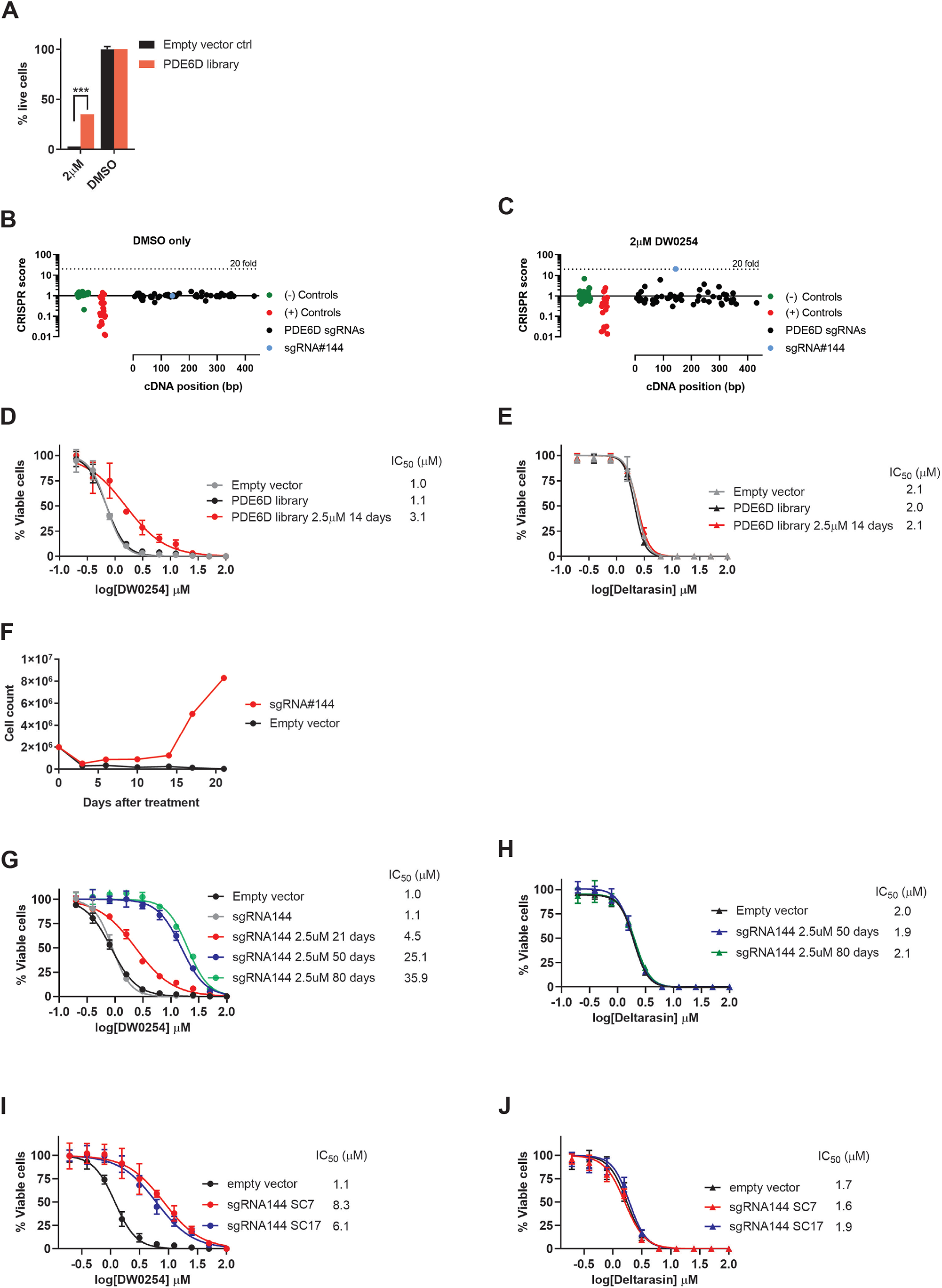
Identification of mutations on V49 and neighboring residues of PDE6D hydrophobic domain as essential for cellular resistance to DW0254. **A)** Percentage of live P12-ICHIKAWA cells by DAPI staining after transduction with either PDE6D library or empty vector control treated for two weeks with 2μM of DW0254 or DMSO, data represent mean ±SD of 3 technical replicates, ***P≤0.001. **B)** Changes in barcoded sgRNAs of untreated PDE6D library cells 14 days after transduction. The cDNA position (in bp) is shown on the X-axis. The fold-change in CRISPR score is shown on the Y-axis. Negative and positive controls are shown in green and red dots, respectively. Negative controls used were non targeting sgRNAs and positive controls targeting essential genes, including PCNA, CDK1, CDK9, RPA3, BRD4, MYC and RPS20. **C)** Changes in barcoded sgRNAs of PDE6D library cells treated for 14 days with 2μM of DW0254 versus 14 days of DMSO. Dotted line on panels B and C represents a 20-fold change on CRISPR score. **D)** DW0254 dose response curves showing % of viable PDE6D library cells or controls untreated or treated with DW0254 at 2.5µM for 14 days. **E)** Deltarasin dose response curves showing % of viable empty vector transduced cells, untreated PDE6D library cells and PDE6D library cells treated with DW0254 for 14 days. **F)** Cell growth curves for P12-ICHIKAWA cells expressing Cas9 only or Cas9 and sgRNA144, treated with 2.5μM of DW0254 for 21 days. **G)** DW0254 dose response curves showing % of viable empty vector transduced cells, untreated sgRNA144 transduced cells, and sgRNA144 cells treated with DW0254 for 21, 50 and 80 days. **H)** Deltarasin dose response curves showing % of viable untreated sgRNA144 transduced cells and controls, and sgRNA144 cells treated with DW0254 for 21, 50 and 80 days; For panels D, E, G and H: data represent mean ± SD of 2 independent experiments with N=3 samples for each condition. **I)** DW0254 dose response curves showing % of viable cells transduced with empty vector and two single cell clones of sgRNA144 transduced cells. **J)** Deltarasin dose response curves showing % of viable empty vector and two single cell clones of sgRNA144 transduced cells; For panels I and J data represent mean ±SD of 2 independent experiments with N=4 samples for each condition.

Next, to define the mutations generated with sgRNA#144 and confirm their association with resistance to DW0254, we transduced spCas9 expressing cells with sgRNA#144 alone. Resulting edited cells demonstrated a resistance phenotype as early as 10 days after treatment, with a 5-fold increase in cell counts when compared to controls, and robust cell growth after day 14 (Figure 4F). Importantly, 100% of empty vector control cells were dead after 17 days of treatment with DW0254 and no resistance was observed in this condition (Figure 4F). Continued selection led to >30-fold increased IC_50_ (Figure 4G). Although highly resistant to DW0254, these edited cells showed no increased resistance to Deltarasin (Figure 4H). Resistant cells genome was enriched for INDELS that would cause the deletion of V49 and neighboring residues within the hydrophobic pocket of PDE6D (data not shown). We validated these predicted mutations using long-range RT-PCR and documented a -6bp in-frame mutation that would cause the combined deletion of R48 and V49 residues (Supplemental Figure 2B) and two out-of-frame mutations (+1bp and -8bp) that both lead to a frame-shift with the formation of a new open reading frame (ORF) with 124 and 127 instead of 150 residues, respectively (Supplemental Figure 2C). Since the new ORF is predicted to translate into a protein that is missing a substantial portion of the hydrophobic pocket and such a change would most likely prevent correct protein folding, we focused our subsequent studies on R48del V49del PDE6D.

To definitively confirm the causal relationship of this mutation to the observed resistance to DW0254, we next isolated sgRNA#144 transduced single cell clones before treatment with DW0254. Edited single cell clones (SC7 and SC17) which harbored R48 and V49 deletions showed a 6-8-fold increased IC_50_ to DW0254 when compared to controls (Figure 4I) while again showing no resistance to Deltarasin (Figure 4J).

### Distinct binding of DW0254 to PDE6D hydrophobic pocket

Next, we determined the binding affinities of the various compounds by isothermal calorimetry (ITC) using recombinant PDE6D protein. For the DW compounds, ITC binding affinity is in line with the order of cellular activities while Deltarasin showed a slightly higher affinity to the protein when compared to DW0254 (Figure 5A). Inactive DW0346 showed very weak affinity with Kd 68.5µM by ITC. Cocrystal structure of DW0254 with recombinant PDE6D shows the small molecule bound inside the hydrophobic pocket, with hydrogen bond interactions via glutamine Q88, tyrosine Y149 and arginine R61, the latter interaction being water mediated (Figure 5B). Deltarasin can occupy the same pocket undergoing hydrogen bonding with the same residues R61 and Y149, but also with cysteine C56 (Figure 5C), which differentiates it from the interactions observed for DW0254. The observed network of hydrogen bonding with the protein backbone supports the strong enthalpy (ΔH) driven binding for both molecules as observed by ITC (Figure 5A).

**Figure 5.**
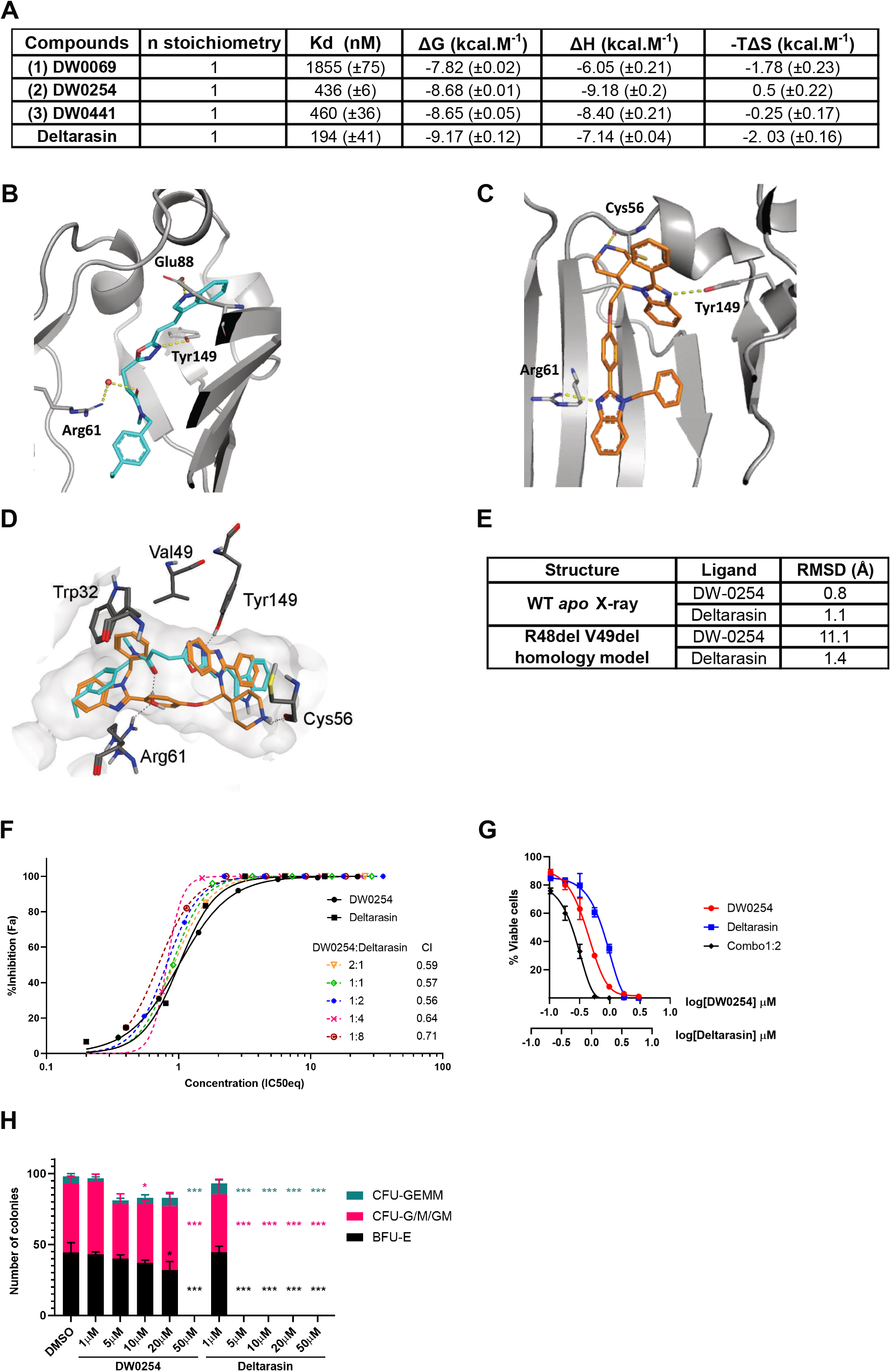
Co-Crystallization of PDE6D and DW0254 confirms compound binding to hydrophobic pocket and evidences different binding modes between this inhibitor and Deltarasin. **A)** Binding affinity (Kd) and thermodynamics parameters for ligand binding to PDE6D determined by Isothermal Titration Calorimetry (ITC). **B)** The crystal structure of compound DW0254 in the PDE6D-binding pocket; Q88, R61 and Y149 are shown in sticks to highlight the hydrogen bind interactions. **C)** The crystal structure of Deltarasin in the PDE6D-binding pocket; C56, R61 and Y149 are shown in sticks to highlight the hydrogen bind interactions. **D)** Experimental binding mode of DW-0254 (cyan) in wild type PDE6D. The superposed 3D coordinates of Deltarasin (orange) are also shown. Several binding site residues, including V49, are shown for reference. **E)** Docking results in wild type apo structure of PDE6D and R48delV49del mutant; Root mean square deviation (RMSD) with respect to 3D coordinates of the ligands in the superposed X-ray complex of the wild type protein are reported. **F)** Drug dosage curves for P12-ICHIKAWA cells treated with DW0254 alone, Deltarasin alone, or various combinations of both obtained from a full matrix using the “Fixed Ratio” Method; data shows mean ± SD of n=3 samples for each condition, and combination indexes for each combo calculated using the Chou-Talalay theorem. **G)** Drug dosage curves for P12-ICHIKAWA cells treated with DW0254 alone, Deltarasin alone, or the combination of both at a 1:2 ratio; data shows mean ± SD of n=3 samples for each condition. **H)** Colony counts of healthy human CD34^+^ cells after 14 days culture in MethoCult H4435 enriched medium in presence of DMSO or increasing concentrations of DW0254 or Deltarasin. CFU-GEMM: Colony-forming unit − granulocyte, erythroid, macrophage, megakaryocyte; CFU-GM: Colony-forming unit − granulocyte, macrophage; BFUE: Burst-forming unit – erythroid. Data represent mean ± SD, n=3 samples for each condition. *p<0.05; ***p<0.001

Guided by the crystallographic information we were also able to postulate a binding pose for the PAL probe (Supplemental Figure 3). To contextualise the crystallographic binding modes with the saturating mutagenesis screen results, superimposing the binding poses of DW0254 and Deltarasin highlighted that V49 defines the shape of the pocket (light grey area, Figure 5D), and establishes hydrophobic contacts only with DW0254 (cyan) but not Deltarasin (orange). Additionally, *in silico* docking of DW0254 to R48del V49del PDE6D confirmed an accentuated increase in the root mean square deviation (RMSD) in contrast with Deltarasin’s RMSD that was only marginally affected (Figure 5E), strongly suggesting DW0254 would be unlikely to bind PDE6D in the event of deletion of these two residues. Interestingly, the combination of DW0254 and Deltarasin had a synergistic effect *in vitro* (Figure 5F) with the lowest combination index at a 1:2 range (Figure 5G), suggesting that even though binding of these compounds to PDE6D is mutually exclusive, they may target different protein conformations more efficiently. However, while DW0254 exhibited low toxicity to CD34^+^ healthy donor cells at therapeutic dosages, Deltarasin showed decreased colony counts even at low dosages (Figure 5H) indicating possible off-target effects of the latter.

### RAS protein dynamics and downstream effects of DW0254

PDE6D has been shown to bind farnesylated RAS proteins and facilitate their trafficking and plasma membrane (PM) localization [24, 25]. To determine the effect of DW0254 on PDE6D-RAS interactions, we generated P12-ICHIKAWA cells that stably expressed a FLAG-tagged human PDE6D protein. Co-immunoprecipitation studies confirmed PDE6D binding to both RAS and ADP-ribosylation factor-like protein 2 (ARL2) protein essential for cargo displacement, that decreased after treatment with DW0254 (figure 6A). No direct binding was observed between PDE6D and RAC (Figure 6A).

**Figure 6.**
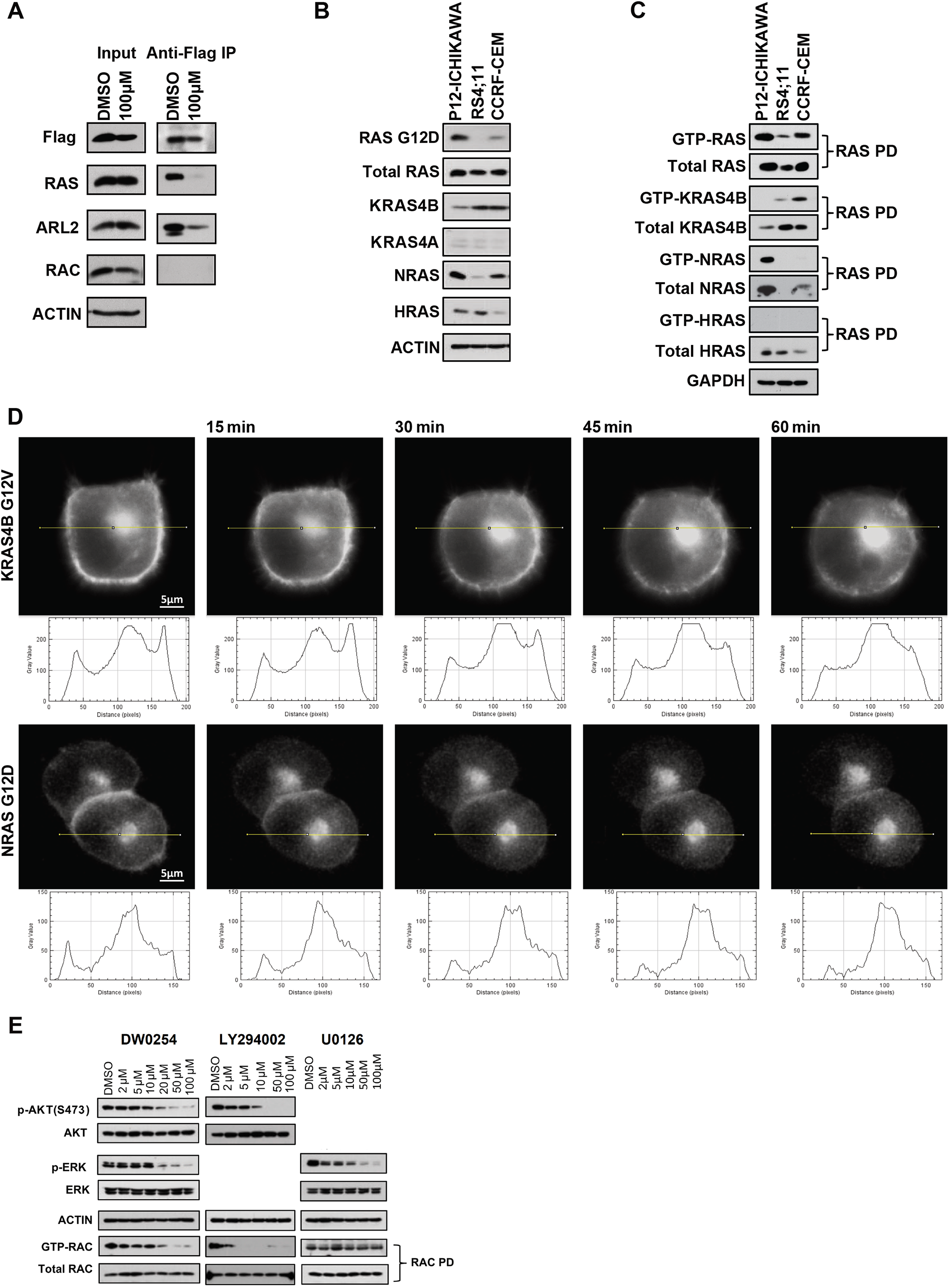
The expression and activation of RAS isoforms in DW0254 sensitive leukemia cells and the effects of DW0254 on PDE6D/RAS interaction and RAS subcellular location. **A)** DW0254 treatment inhibits the binding of PDE6D to RAS and ARL2 in P12-ICHIKAWA cells. Co-immunoprecipitation (CoIP) was performed with an anti-FLAG antibody (F1804, Sigma-Aldrich) on lysates from FLAG-tagged PDE6D transduced P12-ICHIKAWA cells treated with 100µM DW0254 or DMSO. Cell lysates (Input) and protein eluted from beads (IP) were analyzed by Western blotting with anti-Flag, anti-RAS (05-1072, Millipore Sigma, Billerica, MA), anti-ARL2 (ab183510, Abcam, Cambridge, MA), anti-RAC (610651, BD) antibodies. **B)** The expression pattern of RAS isoforms in leukemia cell lines sensitive to DW0254. Western blot analysis of whole-cell lysates from P12-ICHIKAWA, RS4;11, and CCRF-CEM leukemia cells detected by anti-RASG12D (14429S, Cell signaling), anti-RAS (05-1072, Millipore Sigma), Anti-KRAS4B (WH0003845M1, Millipore Sigma), anti-KRAS4A (ABC1442, Millipore Sigma), anti-NRAS (sc-31, Santa Cruz), and anti-HRAS (18295-1-AP, Proteintech, Rosemont, IL) antibodies. **C)** Activated RAS isoforms in P12-ICHIKAWA, CCRF-CEM and RS4;11 cells. GST pulldown assays were performed by incubating protein lysates prepared from P12-ICHIKAWA, CCRF-CEM and RS4;11 with RAF-1 RBD conjugated agarose beads. The GTP-RAS proteins bound to the beads or the whole cell lysates to detect the level of total RAS protein were identified using anti-RAS, anti-KRAS4B, anti-NRAS, or anti-HRAS antibodies described above. For Figure 6A, 6B, and 6C, beta-ACTIN or GAPDH (A300-641A, BETHYL, Montgomery, TX) were used as a protein loading control, one representative experiment of two or three is shown. **D)** Mislocalization from cell surface membrane of GFP-tagged mutant KRAS4BG12V (upper panel) or NRASG12D (lower panel) in PANC-1 cells after treatment with 20µM of DW0254. Time in minutes is indicated above the panels; the first panel represents the moment immediately after the addition of the inhibitor. The intensity profiles show changes in the signal on the Y-axis along the yellow lines on the micrographs above. **E)** Western blot showing total and phosphorylated AKT and ERK, and pulldown results for RAC activation in P12-ICHIKAWA cells treated with increasing doses of DW0254, PI3K inhibitor LY294002, or MEK inhibitor U0126 for 3 hours. Total AKT, and ERK expression were assessed using anti-AKT (9272, Cell signaling) and anti-ERK (9102S, Cell signaling) antibody respectively. Phosphorylation of AKT and ERK were assessed using anti-phospho-AKT Ser473 (9271S, Cell signaling) and anti-phospho-ERK (4377S, Cell signaling) antibody respectively. Total RAC and GTP-RAC were analyzed by RAC pull-down assay as described in Figure 2E. For Figure 6A, 6B, 6C, and 6D, beta-ACTIN or GAPDH were used as a protein loading control. Data represent three independent experiments.

**Figure 7.**
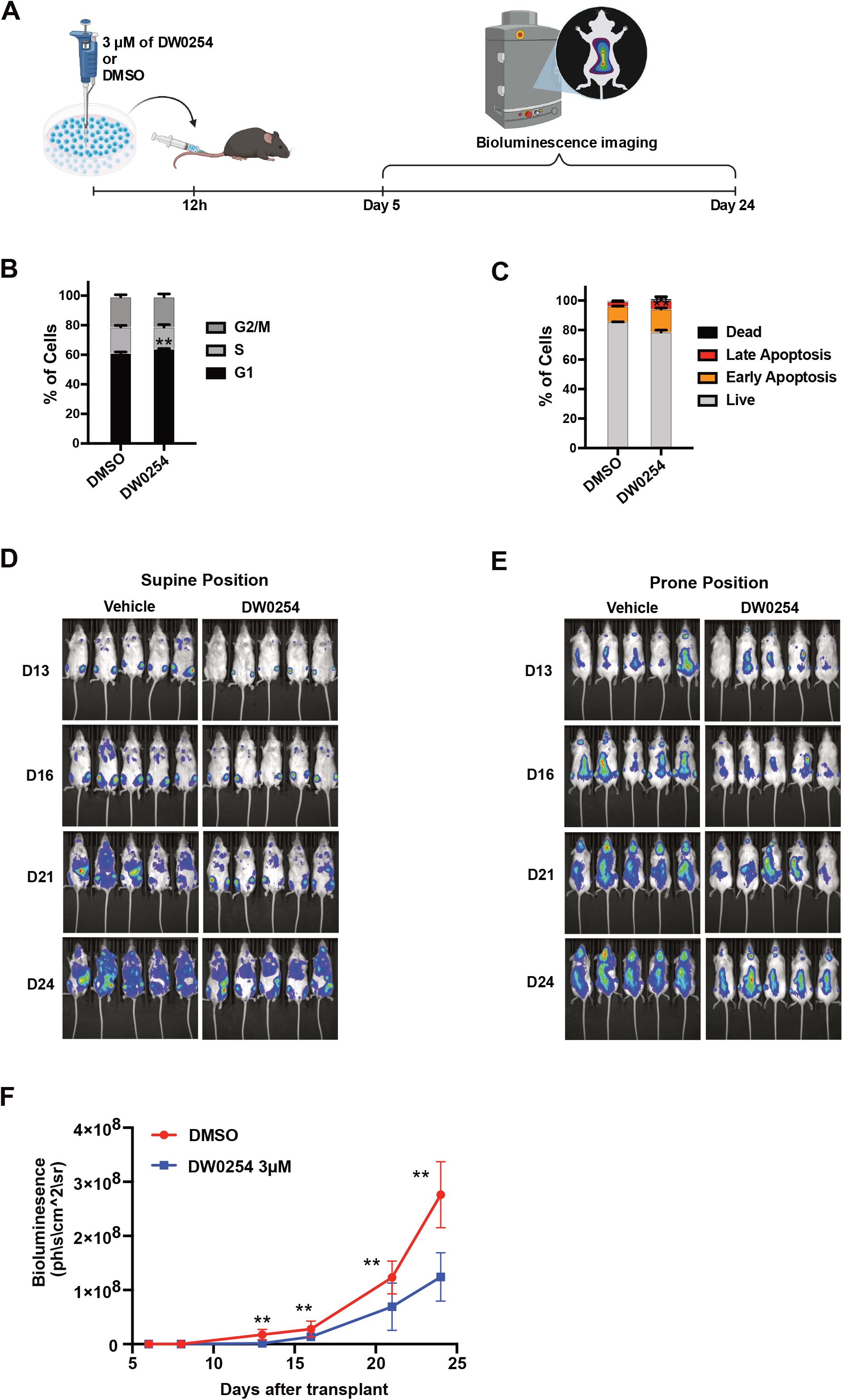
DW0254 *ex vivo* treatment reduces leukemic tumor burden of P12-ICHIKAWA mouse xenograft model. **A)** Schedule of *ex vivo* treatment. Luciferase expressing P12-ICHIKAWA cells treated for 12 hours with either 3μM of DW0254 or DMSO before transplantation into NSG mice; Bioluminescence imaging was performed every 3-5 days to assess tumor burden between day 5 and day 24 after transplantation. Created with BioRender.com. **B)** Bar graph showing the cell cycle distribution by DAPI staining of P12-ICHIKAWA cells treated for 12 hours with 3μM of DW0254, data represent mean ± SD of 2 independent experiments with n=3 samples for each condition, ** p<0.01. **C)** Bar graph showing percentage of apoptosis by AnnV/DAPI staining of P12-ICHIKAWA cells treated for 12 hours with 3μM of DW0254, data represent mean ± SD of 2 independent experiments with n=3 samples for each condition, ** p<0.01. Live: AnnV^-^/DAPI^-^; Early apoptosis: AnnV^+^/DAPI^−^; Late apoptosis: AnnV^+^/DAPI^+^; and Dead: AnnV^−^/DAPI^+^. **D)** Representative images in the prone or **E)** supine position of bioluminescence imaging (BLI) from NSG mice transplanted with P12-ICHIKAWA leukemia cells expressing luciferase after treatment with 3μM DW0254 *ex vivo* for 12 hours. **F)** Quantification of BLI in NSG mice transplanted with luciferase-expressing P12-ICHIKAWA cells after *ex vivo* treatment with 3μM DW0254 for 12 hours. The days after transplantation are shown in X-axis, and the bioluminescence intensity is shown in Y-axis. Data represent mean ±SD, **p<0.01, n=10 mice per group, and were representative of 2 independent experiments.

PDE6D has been reported to chaperone NRAS, HRAS, and KRAS4B but not KRAS4A [25, 26]. While all RAS isoforms were detected in a panel of DW0254-sensitive ALL cell lines, the most abundantly activated isoforms were NRAS in P12-ICHIKAWA and KRAS4B in RS4;11 and CCRF-CEM (Figure 6B and 6C). To further test whether the DW0254-dependent disruption of the interaction between PDE6D and RAS is associated with altered subcellular localization of RAS, we used fluorescently-tagged mutant RAS proteins to analyze RAS localization before and after treatment with DW0254 in adherent PANC-1 cells. Live-cell fluorescence imaging demonstrated that mutant KRAS4B and NRAS dissociated from the PM and accumulated into the cytosol after DW0254 treatment (Figure 6D).

Biological effects of RAS proteins are exerted from the PM through the activation of kinase pathways including PI3K/AKT and MAPK/ERK, which are commonly constitutively activated in cancer [27]. We observed decreased activation of PI3K/AKT and MAPK/ERK pathways, measured by levels of phospho-AKT and phospho-ERK, upon inhibition of PDE6D-RAS interaction by DW0254 (Figure 6E, left panel). Next, to establish a biochemical link between RAC and RAS pathway inhibition we examined if RAC activation is affected by the inhibition of PI3K/AKT or MEK/ERK pathways. The PI3K inhibitor LY294002 clearly decreased GTP-RAC levels while the MEK inhibitor U0126 had no demonstrable effect on RAC activity (Figure 6E). Taken together these results provide a potential molecular link between PDE6D pocket occupation by DW0254, RAS mislocalization, decreased downstream pathway activation, and inhibition of RAC activity.

### DW0254 anti-leukemic activity in a murine xenograft model

Initial *in vivo* pharmacokinetics (PKs) assays demonstrated low solubility and rapid plasma clearance of DW0254 (data not shown) which meant direct *in vivo* treatments could not be performed. We examined the antitumor effects of PDE6D inhibition on leukemia cell growth *in vivo* by treating luciferase-expressing P12-ICHIKAWA cells with DW0254 before transplantation into sub-lethally irradiated non-obese diabetic severe combined immunodeficiency-gamma (NSG) mice (as depicted in Figure 6A). After this short exposure to DW0254 we observed a minimal increase in cells in the G1 phase of the cell cycle (64.1% ± 0.2% for DW0254 vs 61.2% ± 0.7% for DMSO control, p<0.01) and in early apoptosis (15.6% ± 5.7% for DW0254 vs 10.4% ± 3.6% for DMSO control, p<0.01), with negligible effects on late apoptosis and cell death before transplantation (Figure 6B-C). However, disease burden as assessed using bioluminescence imaging was significantly reduced in mice injected with DW0254 treated cells compared to the vehicle control group on days 13, 16, 21 and 24 after injection (Figures 6D-F). In line with the fact that PDE6D inhibition by DW0254 is reversible, *ex vivo* treatment did not result in a survival advantage (data not shown). In summary, the decrease in tumor burden observed up to 24 days after transplant suggests that PDE6D inhibition causes a reduction in the tumorigenic potential of leukemia cells.

## Discussion

The results presented in this manuscript provide evidence of the importance of PDE6D in sustaining RAS activity and consequently, the survival off leukemic cells.

Treatment of ALL cell lines with DW0254 resulted in a clear decrease in GTP RAC. However, binding between DW0254 and RAC was not observed contradicting computer-aided drug design methodologies. We determined the direct target of DW0254 to be PDE6D, a chaperone protein that facilitates cytoplasmic trafficking of farnesylated molecules, including RAS, as a target for this compound, thus linking RAS transport with Rac GTPase activation in leukemic cells. Saturating mutagenesis experiments showed that the deletion of R48 and V49 residues causes changes to PDE6D pocket that prevent binding to DW0254 and result in resistance to the compound. The binding mode for DW0254 in PDE6D farnesyl binding pocket was also confirmed by crystallography and is different than the binding mode of another previously described inhibitor, Deltarasin. Further emphasizing the importance of this difference, R48del V49del edited cells are not resistant to Deltarasin. Interestingly, the combination of DW0254 and Deltarasin had a significant synergistic effect suggesting that these compounds might be targeting singular conformations of PDE6D with different efficiencies. Indeed, large conformational changes in PDE6D to facilitate the binding of farnesylated RAS proteins deeper within the hydrophobic pocket have been previously described [28]. Additionally, DW0254 did not show any toxicity to CD34^+^ healthy donor cells at therapeutic levels, suggesting a potential for translational improvement of this inhibitor. Even though a role for PDE6D on blood cell differentiation has not been previously described, low dosages of Deltarasin led to decreased colony counts.

DW0254 treatment leads to the delocalization of RAS from the plasma membrane, where it can activate downstream factors [29], to the cytoplasm, as had been previously reported with other PDE6D inhibitors [23]. As shown here and in line with recent studies on the importance of RAS membrane localization [29], RAS delocalization ultimately results in an inability to activate target pathways including MAPK/ERK, PI3K/AKT, and consequently RAC.

Active KRAS4B, which is solely dependent on PDE6D trafficking for its transport to the membrane, was found in WT RAS cell line RS4;11 which is highly responsive to DW0254. As discussed, delocalization of KRAS4B downstream of PDE6D pocket occupation by DW0254 leads to PI3K/AKT inhibition which has previously been implicated in RS4;11 cell death [30]. Together with the fact that IC_50_ values did not correlate with RAS mutational status of acute leukemia cell lines, this suggests that RAS pathway activation might be a better predictor of response to PDE6D inhibition. However, one additional target of DW0254 was identified in one ALL cell line by PALMS assay with a lower signal intensity (Log2 Intensity 20.50 compared with 24.66 for PDE6D) and sequence coverage (8.1% compared to 28.6% for PDE6D) (data not shown). This target is being validated and could also contribute to the effects observed on leukemic cell division and viability.

In conclusion, we have validated the RAS chaperone PDE6D as a novel molecular target for aggressive leukemias. We have derived a series of compounds with demonstrated PDE6D inhibition that bind to its hydrophobic pocket differently from a previously identified inhibitor series showing little toxicity to normal human and mouse hematopoietic progenitor cells. The binding of DW0254 to PDE6D resulted in delocalization of RAS from the membrane and consequent inhibition of major pro-survival pathways including MAPK/ERK, PI3K/AKT and downstream RAC activation. These results also suggest that combinatorial strategies that inhibit parallel signaling through these pathways may increase the anti-leukemic responses and become particularly clinically significant in treating relapsed patients.

## Methods

### Cell lines

CCRF-CEM, RS4;11, MV4;11, and PANC-1 cells were obtained from ATCC and all others from DSMZ. Cells were cultured according to suppliers’ instructions and periodically tested for the presence of mycoplasma.

### Cell Viability Assay

Cells were treated for 3 days at 1×10^5^ cells/ml with limiting dilutions of DW0254 or DMSO only. On day 3, cells were stained with DAPI at a 1 μg/ml final concentration and the number of viable (DAPI-) cells in 25μl of media were counted using BD LSR II.

### AnnV/PI Staining and Cell Cycle Analysis

P12-ICHIKAWA cells were plated at a 2×10^5^ cells/ml concentration with DW0254 or DMSO for 3 days. Cells were labelled with Dead Cell Apoptosis Kit with Annexin V FITC and PI (Thermo Fisher) or fixed in 70% ethanol at 4°C overnight, followed by incubation with 10µg/mL Ribonuclease A (Sigma-Aldrich, St Louis, MO) and 50µg/mL PI (BD Biosciences PharMingen, San Diego, USA) or 10µg/mL DAPI (Thermo Fisher). Flow cytometry analysis was performed on a BD LSR II.

### Generational Cell Tracing

Cells were stained with CellTrace™ Far Red (Thermo Fisher) following manufacturer’s instructions and incubated with DW0254 or DMSO. Cells were analyzed on BD LSR II after 15 minutes (Time 0) and the following 3 days at the same time.

### Recombinant Protein Expression and Purification

Recombinant human Rac1 (Q2-L177) with TEV-protease cleavable 6xHis-tag fused to its N-terminus, and truncated recombinant human Tiam1 (R1033-E1406) with FLAG-tag fused to its N-terminus were cloned into in the pTriIJ-HV vector and expressed in BL21 (DE3). Rac1 protein went through a nickel affinity column followed by a Resource Q column and finally Superdex 75 (GE Healthcare) before concentration to 25mg/ml. Tiam1 protein was purified using the ANTI-FLAG® M2 affinity gel (Sigma-Aldrich) followed by Superdex 75. Recombinant human PDE6D (S2-V150) with TEV-protease cleavable 6xHis-tag fused to its N-terminus, was cloned into pET28a, expressed in BL21-CodonPlus (DE3)-RIL and purified using nickel affinity chromatography followed by TEV protease cleavage, tag removal, and finally Superdex 75 before concentration to 13mg/ml.

### Isothermal calorimetry (ITC)

PDE6D was dialyzed in buffer (20mM HEPES pH7.3, 150mM NaCl, 1mM TCEP) at 4°C, overnight. Titrations were carried out on an iTC200 calorimeter (MicroCal Inc). PDE6D (200µM with 2% DMSO) was titrated into small molecule in the cell (20µM in degassed dialysis buffer with 2% DMSO final) and data were analyzed using Origin (OriginLab Corp.) and fitted by using a single-site binding model.

### Rac1-Tiam1 homogeneous time-resolved fluorescence assay (HTRF)

30nM His-tagged Rac1 protein was pre-incubated with compound at room temperature in assay buffer (50mM Hepes (pH 7.6), 100mM NaCl, 1mM DTT, 10mM MgCl_2_, 0.1% Nonidet P-40). After 30 minutes pre-incubation, 300nM FLAG-tagged Tiam1, 2nM anti-His-Eu3+, 20nM anti-FLAG-XL665 were added. After 60 minutes RT incubation, 500mM Potassium Fluoride (KF) was added and the reaction was measured after 30 minutes with EnVision 2104 Multilabel Reader (Perkin Elmer) with the following settings. Ex: 320nm; Em1: 615nm; Em2: 665nm; Dichroic Mirror: D400.

### High Density sgRNA Library of Human PDE6D

sgRNA sequences targeting the coding regions of human PDE6D (NM_002601.3) were designed using Genetic Perturbation Platform from Broad Institute [31] (Supplemental table 1). Briefly, sgRNA oligonucleotides were synthesized via microarray (CustomArray) and cloned into the ipUSEPR lentiviral sgRNA vector that co-expresses a puromycin-resistant gene [puro^R^] and a red fluorescent protein [tagRFP]. The PDE6D scan library contains 116 unique sgRNA was packaged by HEK293 cells (ATCC) co-transfected with psPAX2 (Addgene) and pMD2.G (Addgene) to produce lentiviral particles. The lentiviral library was pre-titrated to obtain 5-10% infection (monitored by flow cytometry for tagRFP expression from ipUSEPR) in P12-ICHIKAWA spCas9 expressing cells. Each screen culture was calculated to maintain at least 1,000x of the number of constructs in the library. The infected cultures were selected by sorting of RFP^+^ cells 3 days after transduction and expanded in supplemented media with puromycin (2.5µg/ml; InvivoGen) and blasticidin (1µg/ml; InvivoGen) for 3 additional days. Finally, selected cells were pelleted (day 0) and cultured in DMSO or 2.0µM DW0245. After 14 days treatment cells were again pelleted. For sequencing sgRNAs, the genomic DNA of the screened cell pellets was harvested, PCR-amplified (NEBNext Ultra II Q5; NEB) using primers DCF01 5’-CTTGTGGAAAGGACGAAACACCG-3’ and DCR03 5’-CCTAGGAACAGCGGTTTAAAAAAGC-3’ and subjected to single-end 75 bp (SE75) high-throughput sequencing using a NextSeq550 (Illumina).

To quantify sgRNA reads in the library, we first extracted 20-nucleotide sequences that matched the sgRNA backbone structure (5’ prime CACCG and 3’ prime GTTT) from raw fastq reads. Extracted reads were then mapped to the PDE6D sgRNA library sequences using Bowtie2 [32]. Reads that were a perfect match to the reference were counted. The frequency for individual sgRNAs was calculated by the read counts of each sgRNA divided by the total read counts matched to the library. The CRISPR score was defined by the fold change of the frequency of individual sgRNAs between early (day 0) and late (defined time points) of the screened samples.

### Crystallization and Structural Determination

Native PDE6D crystals were grown by vapor diffusion at 22°C by mixing equal volumes of protein with precipitant (0.1M HEPES pH6.8-7.5, 20mM MgCl_2_, 20mM NiCl_2_ and 15-20% PEG3350). DW0254 and Deltarasin were incubated with PDE6D at 4°C, at 4mM and 1mM final respectively. The PDE6D::Deltarasin complex was further concentrated to 19 mg/ml prior setting up the crystallization trays. PDE6D::Deltarasin and PDE6D::DW0254 complexes were grown by vapor diffusion at 22°C in (0.1 M Sodium acetate pH4.0-4.5 and 28−30% PEG3350) and (0.1M HEPES pH6.8-7.5, 20mM MgCl_2_, 20mM NiCl_2_ and 15-20% PEG3350) respectively. Prior to freezing in liquid nitrogen, crystals were cryoprotected by brief transfer to a solution of crystallization condition reservoir supplemented with 25% glycerol. Data were collected on beamlines at Diamond Light Source (Oxford, U.K.) and ALBA (Barcelona, Spain). Data were processed using XDS, xia or DIALS. Molecular replacement was performed using PHASER (using PDB code 5NAL as a reference model), and the refinement was performed with refmac5, buster and COOT. Compound dictionaries were generated using AFITT.

### Combination Index Analysis

Each drug was used alone or in combination at a concentration approximately equal to its IC50 and at concentrations within 2-2.5-fold increments above or below. Each data point was performed in triplicates. In this model, combination index (CI) scores estimate the interaction between the two drugs. If CI<1, the drugs have a synergistic effect [33]. To allow a direct comparison of the dose-response curves, each drug concentration was normalized to its own IC_50_ value and named IC_50_ equivalent (IC50eq) as previously described by Zhao et.al. [34]:

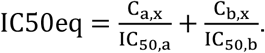

### PDE6D Co-immunoprecipitation (Co-IP)

NH3-terminal FLAG-tagged human PDE6D was constructed by PCR, checked by sequencing, and subcloned into the BglII and EcoRI site of MSCV-IRES-GFP vector. GFP^+^ P12-ICHIKAWA cells were sorted 48 hours after lentiviral infection.

Cells with stable expression of FLAG-tagged human PDE6D were lysed in 1X cell lysis buffer (#9803, Cell Signaling, Danvers, MA) and incubated with anti-FLAG M2 Affinity Gel (A2220, Sigma-Aldrich) overnight at 4°C. Protein complexes were washed 5 times with 1mL lysis buffer, then 2X SDS sample buffer was added, following 100°C incubation for 5min.

### RAS and RAC activity assay

RAS and RAC activity were measured using a RAF-1 RBD and PAK-1 PDB pull-down assay kits respectively (Cat#17218 and Cat#14325, Millipore Sigma) following manufacturer’s instructions. For comparison with total corresponding protein, 5–10% of total lysates used for pulldown were loaded to adjacent wells.

### Transfection and Fluorescent Imaging

PANC-1 cells were collected from a confluent flask, split 1:5 and plated on 35mM µ-Dishes with a polymer coverslip bottom (Ibidi) and incubated in a humidified 37°C incubator with 5% CO_2_ for 24 hours. The next day cells were transfected with pEGFP-C3 KRAS4B 12V or pEGFP-C3 NRAS 12D using Lipofectamine 3000 (Thermo Fisher) following manufacturer’s instructions, and incubated for 3 days in a humidified 37°C incubator with 5% CO_2_. Cells in 1.8ml PBS with 10%FCS were imaged in a Nikon Eclipse Ti inverted microscope with a humidified live cell imaging chamber using NIS-Elements software. 200µl of PBS with 10% DMSO only or of 200µM DW0254 previously diluted in PBS with 10% DMSO were added, and samples were imaged every 5 minutes for 1 hour.

### Bioluminescent imaging for DW0254 *ex vivo* efficacy studies

To generate a cell line with luciferase expression, P12-ICHIKAWA cells were infected with Lenti-FUW-Luc-mCh-puro virus and selected in liquid culture with puromycin (Sigma-Aldrich) 2.5µg/mL for 7 days following mCherry^+^ cell sorting.

All animal studies were approved by the Boston Children’s Hospital or Dana-Farber Cancer Institute Animal Care and Use Committee. 8- to 10-week-old NOD-scid IL2Rgamma^null^ (NSG) mice (Jackson laboratories, Bar Harbor, ME) were sublethally irradiated with 300 cGy and injected with 1×10^6^ luciferase expressing P12-ICHIKAWA cells treated for 12 hours with DMSO or 3µM DW0254. Disease burden was assessed using bioluminescence imaging starting six days after injections. Prior to imaging, each mouse was given an intra-peritoneal (i.p.) injection of luciferin (PerkinElmer, Part Number #122799) at a dose of 150mg/kg body weight. General anesthesia was then induced with 5% isoflurane and the mouse was placed in the light-tight heated chamber; anesthesia was continued during the procedure with 2% isoflurane introduced via nose cone. Both prone and supine images were recorded.

Optical images were displayed and analyzed with the Igor (WaveMetrics, Lake Oswego, OR) and IVIS Living Image (Xenogen) software packages. Regions were manually drawn around the bodies of the mice to assess signal intensity emitted. Optical signal was expressed as photon flux, in units of photons/s/cm^2^/steradian. The total photon flux for each mouse was calculated as the sum of prone and supine photon flux.

### PAL Probe Synthesis, Photoaffinity Labelling and LC-MS/MS

All the information regarding the synthesis of PAL probe, and specifics on photoaffinity labelling and LC-MS/MS data collection and analysis are available under supplementary information.

### Statistical analysis

Data were presented as mean ±SD. The unpaired t test was used for comparisons between groups at each time point. P<.05 was considered significant.

## Data Availability

The coordinates for the apo PDE6D alone and bound to Deltarasin or DW0254 have been deposited in the PDB under accession codes 7PAC, 7PAE and 7PAD respectively. Authors will release the atomic coordinates and experimental data upon article publication.

## Acknowledgments

The authors thank the Flow Lab HSCI Core at BCH for their help in cell sorting experiments; Hiroko Hishikawa from the BCH ARCH team for help with IVIS setup; Mark Philips for the mutant Ras plasmids; Jenna Wood for animal husbandry and experimental support; Mursal Hassan and Timothy Colby for assistance in manuscript preparation and submission; Alejandro Gutierrez, Scott Armstrong, Nathanael Gray, and the members of the Williams laboratory for the helpful discussions.

This work is supported by grant 5R01CA202756 (D.A.W.), and a ALSF Young Investigator Award 19-16300 co-funded by Alex’s Lemonade Stand Foundation and Cure4Cam (S.C.N.).

## Author contributions

S.C.N., S.D.V., A.A., F.A., E.B., P.K., M. MG., B. D., C. H., H.X. conducted experiments and/or data analysis. S.C.N., F.A., E.B., CW. C., M.E., D.A.W. and H.X. designed experiments. S.C.N., F.A., E.B., M.E., D.A.W. and H.X. wrote the paper.

## References

1. Troeger, A. and D.A. Williams, Hematopoietic-specific Rho GTPases Rac2 and RhoH and human blood disorders. Exp Cell Res, 2013. 319(15): p. 2375–83.

2. Cancelas, J.A., et al., Rac GTPases differentially integrate signals regulating hematopoietic stem cell localization. Nat Med, 2005. 11(8): p. 886–91.

3. Bos, J.L., ras oncogenes in human cancer: a review. Cancer Res, 1989. 49(17): p. 4682–9.

4. Prior, I.A., P.D. Lewis, and C. Mattos, A comprehensive survey of Ras mutations in cancer. Cancer Res, 2012. 72(10): p. 2457–67.

5. Tyner, J.W., et al., High-throughput sequencing screen reveals novel, transforming RAS mutations in myeloid leukemia patients. Blood, 2009. 113(8): p. 1749–55.

6. Neri, A., et al., Analysis of ras oncogene mutations in human lymphoid malignancies. Proc Natl Acad Sci USA, 1988. 85: p. 9268–9272.

7. Loh, M.L., Recent advances in the pathogenesis and treatment of juvenile myelomonocytic leukaemia. Br J Haematol, 2011. 152(6): p. 677–87.

8. Zhang, J., et al., The genetic basis of early T-cell precursor acute lymphoblastic leukaemia. Nature, 2012. 481(7380): p. 157–63.

9. Perentesis, J.P., et al., RAS oncogene mutations and outcome of therapy for childhood acute lymphoblastic leukemia. Leukemia, 2004. 18(4): p. 685–92.

10. Irving, J., et al., Ras pathway mutations are prevalent in relapsed childhood acute lymphoblastic leukemia and confer sensitivity to MEK inhibition. Blood, 2014. 124(23): p. 3420–30.

11. Oshima, K., et al., Mutational landscape, clonal evolution patterns, and role of RAS mutations in relapsed acute lymphoblastic leukemia. Proc Natl Acad Sci U S A, 2016. 113(40): p. 11306–11311.

12. Canon, J., et al., The clinical KRAS(G12C) inhibitor AMG 510 drives anti-tumour immunity. Nature, 2019. 575(7781): p. 217–223.

13. Hallin, J., et al., The KRAS(G12C) Inhibitor MRTX849 Provides Insight toward Therapeutic Susceptibility of KRAS-Mutant Cancers in Mouse Models and Patients. Cancer Discov, 2020. 10(1): p. 54–71.

14. Qiu, R.G., et al., An essential role for Rac in Ras transformation. Nature, 1995. 374(6521): p. 457–9.

15. Thomas, E.K., et al., Rac guanosine triphosphatases represent integrating molecular therapeutic targets for BCR-ABL-induced myeloproliferative disease. Cancer Cell, 2007. 12(5): p. 467–78.

16. Mizukawa, B., et al., Inhibition of Rac GTPase signaling and downstream pro-survival Bcl-2 proteins as combination targeted therapy in MLL-AF9 leukemia. Blood, 2011.

17. Cox, A.D., et al., Drugging the undruggable RAS: Mission possible? Nat Rev Drug Discov, 2014. 13(11): p. 828–51.

18. Williams, D.A., Compounds for treating Rac-GTPase mediated disorder. Patent Int. WO, 2014. 059305.

19. Gao, Y., et al., Rational design and characterization of a Rac GTPase-specific small molecule inhibitor. Proc Natl Acad Sci U S A, 2004. 101(20): p. 7618–23.

20. Wei, J., et al., Microenvironment determines lineage fate in a human model of MLL-AF9 leukemia. Cancer Cell, 2008. 13(6): p. 483–95.

21. Levay, M., et al., NSC23766, a widely used inhibitor of Rac1 activation, additionally acts as a competitive antagonist at muscarinic acetylcholine receptors. J Pharmacol Exp Ther, 2013. 347(1): p. 69–79.

22. Li, Z., et al., Design and synthesis of minimalist terminal alkyne-containing diazirine photo-crosslinkers and their incorporation into kinase inhibitors for cell-and tissue-based proteome profiling. Angew Chem Int Ed Engl, 2013. 52(33): p. 8551–6.

23. Zimmermann, G., et al., Small molecule inhibition of the KRAS-PDEdelta interaction impairs oncogenic KRAS signalling. Nature, 2013. 497(7451): p. 638–42.

24. Chandra, A., et al., The GDI-like solubilizing factor PDEδ sustains the spatial organization and signalling of Ras family proteins. Nat Cell Biol, 2011. 14(2): p. 148–58.

25. Tsai, F.D., et al., K-Ras4A splice variant is widely expressed in cancer and uses a hybrid membrane-targeting motif. Proc Natl Acad Sci U S A, 2015. 112(3): p. 779–84.

26. Chandra, A., et al., The GDI-like solubilizing factor PDEdelta sustains the spatial organization and signalling of Ras family proteins. Nat Cell Biol, 2011. 14(2): p. 148–58.

27. Rajalingam, K., et al., Ras oncogenes and their downstream targets. Biochim Biophys Acta, 2007. 1773(8): p. 1177–95.

28. Dharmaiah, S., et al., Structural basis of recognition of farnesylated and methylated KRAS4b by PDEdelta. Proc Natl Acad Sci U S A, 2016. 113(44): p. E6766–E6775.

29. Tan, L., et al., An oxanthroquinone derivative that disrupts RAS plasma membrane localization inhibits cancer cell growth. J Biol Chem, 2018. 293(35): p. 13696–13706.

30. Gorlick, R., et al., Testing of the Akt/PKB inhibitor MK-2206 by the Pediatric Preclinical Testing Program. Pediatr Blood Cancer, 2012. 59(3): p. 518–24.

31. Doench, J.G., et al., Optimized sgRNA design to maximize activity and minimize off-target effects of CRISPR-Cas9. Nat Biotechnol, 2016. 34(2): p. 184–191.

32. Langmead, B., et al., Ultrafast and memory-efficient alignment of short DNA sequences to the human genome. Genome Biol, 2009. 10(3): p. R25.

33. Chou, T.C. and P. Talalay, Quantitative analysis of dose-effect relationships: the combined effects of multiple drugs or enzyme inhibitors. Adv Enzyme Regul, 1984. 22: p. 27–55.

34. Zhao, J., K. Kelnar, and A.G. Bader, In-depth analysis shows synergy between erlotinib and miR-34a. PLoS One, 2014. 9(2): p. e89105.

